# Central carbon metabolism switching in lytic versus temperate coral reef viral communities

**DOI:** 10.1101/2025.09.25.676695

**Authors:** Jacob Kelman, Meena Khan, Chibundu Umunna, Russell Brainard, Grace Donohue, Rob Edwards, Natalie A Falta, Emma George, Eleanor Gorham, Juris Grasis, Kevin Green, Andreas Haas, Kimberly Halsey, Eric Hester, Summer Jacob, Aydin Loid Karatas, Yan Wei Lim, Mark Little, Stuart Sandin, Jessie Segnitz, Maya Serota, Natalia Shahwan, Giselle Simmons, Jennifer E Smith, Isha Tripathi, Linda Wegley Kelly, Lauren Woodward, Nickie Yang, Charles Young, Brian Zgliczynski, Forest Rohwer, Ben Knowles

## Abstract

Coral reefs are declining globally due in part to bacterial overgrowth, a process known as microbialization. However, the role of bacteriophages that may inhibit microbialization by infecting and killing these bacteria remains poorly understood, especially their metabolic impacts on bacterial proliferation. To address this, we analyzed central carbon metabolism gene frequencies in viral communities from healthy (lytic-dominated) and degraded (temperate-dominated) Central Pacific coral reefs. We found that viral communities shifted broadly from enrichment in genes associated with reactions that build up pools of central intermediates on degraded, temperate-dominated reefs (‘anaplerotic’ reactions) to enrichment in genes associated with reactions that consume these pools to generate metabolic precursors for virion construction on healthy, lytic-dominated reefs (‘cataplerotic’ reactions). This switch was shown by the over-representation of Entner-Doudoroff (ED) glycolysis genes on degraded, temperate-dominated reefs and, on healthy, lytic-dominated reefs, by a cross-pathway enrichment of cataplerotic genes, including transaldolase and NADH-producing oxidoreductases. To explore the ecological implications of these observations, we incorporated them into a conceptual model with coral, algae, bacterial, and viral ‘populations’ and qualitative ‘high’ and ‘low’ rates linking them based on our findings and existing knowledge of healthy and degraded reefs. The model suggests two contrasting ecosystem scenarios: one in which lytic viral metabolism may enhance viral production and bacterial control, and another in which temperate viral metabolism may favor bacterial proliferation. This framework identifies viral metabolism as an important and underexplored dimension of coral reef ecology and a promising avenue for future conservation research.

**Significance:** Coral reefs are collapsing worldwide, exacerbated by ‘microbialization,’ where algae-fueled bacteria overgrow corals. Viruses that infect these bacteria can suppress this process through lysis, but on degraded reefs they often switch to nonlethal temperate lifestyles, accelerating decline. Here we show that similar shifts occur in virus-associated metabolic gene content. On healthy reefs, lytic-associated viral communities contained higher frequencies of genes known to drain host metabolites to fuel virus production. At the ecosystem level, this may enhance viral production, infection and lysis rates, thereby helping to limit bacterial overgrowth. On degraded reefs, temperate-associated viral communities contained higher frequencies of genes whose products are known to expand host metabolite pools, possibly providing the energetic substrate to support bacterial proliferation. Taken together, our findings allow us to propose a framework linking viral infection strategy, viral metabolism, and reef microbialization. Within this framework, viral metabolism may either reinforce reef resilience or exacerbate collapse, providing a foundation to guide future work on the role of viruses in coral reef function and conservation.

## Introduction

Ecosystem overgrowth by microbes, a process known as microbialization, is a major threat to coral reefs worldwide^1–3^. Microbialized reefs are subject to a positive feedback loop in which proliferating algae generate excess dissolved organic carbon (i.e., sugar) that fuels the growth of increasingly pathogenic and copiotrophic microbes that thrive in nutrient-rich environments^1,4,5^. This process exacerbates coral diseases and clears space for further algal growth^3,6–8^.

Bacterial metabolism on degraded coral reefs is directed towards high-throughput rather than efficient carbohydrate utilization^1,9^. For instance, while bacterial metabolism on healthy reefs is dominated by genes for the efficient but multi-step Embden-Meyerhof-Parnas (EMP) pathway, degraded reefs exhibit greater representation of genes for the inefficient but short Entner-Doudoroff (ED) glycolysis pathway^1,10–12^. This shift on degraded reefs enables bacteria to readily exploit ecosystem organic carbon enrichment to build up cellular stockpiles of central metabolites like glyceraldehyde-3-phosphate (G3P) and pyruvate (‘anaplerotic’ reactions) as well as use those central metabolite pools to produce cellular energy such as ATP, reductants like NADH, and amino and nucleic acid biosynthetic precursors required for bacterial over-proliferation (‘cataplerotic’ reactions)^1,7,10^.

Viral predation could, in principle, preclude microbialization by killing proliferating microbes, as lytic infection and lysis rates are predicted to rise with host densities and growth rates by established predator-prey models^13–17^. However, viral communities switch from lytic to temperate infection during eutrophication and microbialization^14^. This transition towards non-lethal, temperate infection can facilitate further increases in microbial biomass due to suppressed lysis^14,17^. Further, temperate viral communities encode higher frequencies of pathogenicity genes that could protect bacteria from consumption by eukaryotic grazers^14,18–20^. These genes could facilitate microbialization by limiting protist predation, the major non-viral predation guild in the ocean^14,20–22^. Altogether, rather than controlling bacterial populations, viruses may form a second ‘Temperate Ratchet’ feedback loop that traps ecosystems in persistent microbialization through suppressed lysis rates and protist grazing on pathogenic copiotrophs^14,23^.

In addition to these population-level effects, viruses may also exert gene-level effects on microbialization by reshaping bacterial metabolism. Viruses shuttle a broad array of metabolic genes between hosts via horizontal gene transfer, thereby altering bacterial metabolism and ecology^24–27^. These include central carbon metabolism genes implicated in the metabolic switching of bacterial communities on microbialized reefs, such as the pentose phosphate pathway (PPP) and the ED and EMP glycolysis pathways^1,28^. In lytic viruses, these pathways support virion synthesis by converting general cellular metabolic pools like G3P and pyruvate to specialized metabolic precursors (i.e., they are cataplerotic reactions). For example, the PPP provides nucleic acid and amino acid precursors, whereas EMP generates cellular energy like ATP and NADH^25,29,30^. However, these genes can also be co-opted by bacteria during infection. As a result, while viral metabolic genes could contribute to suppressing microbialization by promoting the production of lytic viruses that would suppress bacterial abundance, they are also capable of exacerbating reef decline by enhancing bacterial proliferation. Together, these processes introduce a novel viral-mediated metabolic mechanism of reef decline or resilience beyond lytic-to-lysogenic switching.

However, the role of viral-encoded metabolic genes in coral reef microbialization remains unexplored. To address this gap, we leveraged 19 metagenomes of virus-like particles (VLPs; viromes) used in several prior publications^14,31,32^ from Pacific coral reefs spanning gradients of reef condition and viral community temperateness. We characterized changes in the frequencies of viral metabolic genes across this lytic-to-temperate spectrum. Doing so, we found that temperate viral communities were enriched in anaplerotic central carbon metabolism genes, particularly those associated with Entner-Doudoroff glycolysis and the generation of central metabolite pools for diverse cellular processes^33^. In contrast, lytic viral communities were enriched in cataplerotic reactions, including genes from the non-oxidative pentose phosphate pathway and oxidoreductases that generate metabolic precursors, cellular energy, and redox potential.

These metabolic changes add to already complicated feedback loops among viruses, bacteria, corals, and algae involved in reef resilience and degradation. To better understand whether these viral metabolic profiles might reinforce or oppose these established ecosystem processes, we developed a conceptual model that incorporated viral metabolism into this broader ecological context and simulated the resulting system dynamics. Although the model is best used as a conceptual guide to understanding the ecological implications of our findings, it also supports hypothesis generation by suggesting that lytic viral metabolism may enhance viral production and microbial surveillance on healthy reefs (‘viralization’)^34^, while temperate viral metabolism may accelerate microbialization in degraded systems.

Together, our observations and the resulting conceptual framework suggest that shifts between lytic- and temperate-associated metabolism may rebalance existing ecological feedback loops, contributing to the divergent reef fates of either resilience or microbialization. Because switching between lytic and temperate infection is strongly influenced by host physiology and growth state^35^, this framework predicts that interventions favoring lytic infection, for example by limiting nutrient inputs, may influence reef resilience through their effects on viral infection strategy and metabolism.

## Methods

### Metagenome sampling and processing

Our work leverages 19 metagenomes of virus-like particles (VLPs; viromes), all but two of which have been extensively used in prior publications (see **Supplementary Table 1** for sample details)^14,31,32^. These metagenomes were sequenced from virus-like particles sampled from coral reefs across the central Pacific Ocean and processed according to established molecular biology pipelines described previously^14,36,37^. For each viral metagenome, approximately 60-100 L of seawater were pumped from the reef surface into acid-washed, triple-rinsed low-density polyethylene bags and concentrated to < 500 mL using a 100 kDa tangential flow filter. Concentrates were then 0.45 μm-filtered to remove bacteria and 0.5 % final concentration chloroform was added to destroy any residual cells and extracellular vesicles^14,36,37^. Samples were stored at 4 °C until subsequent processing. The presence of concentrated VLPs was confirmed by epifluorescence microscopy of Sybr Gold stained viruses on 0.02 μm anodisc filters^37^.

VLPs were purified from concentrates using a cesium chloride step gradient approach^14,36,37^. The absence of cells in step gradient outputs was confirmed by microscopy. Purified samples were then DNAse I-treated at 37 °C for 2 hours and denatured at 65 °C for 15 minutes to remove extra-virion free DNA (peDNA; protected environmental DNA)^38^. Viral DNA was extracted using the formamide/chloroform isoamyl alcohol technique^14,36,37^. The absence of contaminating host DNA in all samples was confirmed by the lack of PCR amplification of the universal 16S rDNA gene using primers 27F (AGA GTT TGA TCC TGG CTC AG) and 1492R (TAC GGY TAC CTT GTT ACG ACT T)^14,36,37^. VLP DNA was amplified using the Linker Amplified Sequencing Library approach^39^ prior to sequencing on an Illumina MiSeq platform.

### Bioinformatic processing

Low-quality reads < 100 bp in length and with mean PHRED quality scores of < 25 were excluded using Prinseq^40^. Samples were then dereplicated with TagCleaner^41^, human contaminants were removed using DeconSeq^42^, and sequences were posted to MG-RAST (www.mg-rast.org; see Data Availability section). Accession numbers and summary statistics for the viral metagenomes used in this paper are shown in **Supplementary Table 1**.

### Annotating metagenomes

Because this study builds directly on previously published analyses of these viromes^14,31,32^, we retained a read-based analytical framework broadly similar to that used in these prior papers while extending the analysis using the MG-RAST annotation platform to ensure methodological continuity and direct comparability with those studies while extending the analysis to viral metabolic gene repertoires. To this end, reads from viral metagenomes were annotated by DIAMOND alignment against the SEED subsystems database on MG-RAST v4 using default MG-RAST parameters^43^: 10^-5^ e-value, 60 % identity, a minimum alignment length of 15 amino acids, a minimum abundance of 1, and all analyses conducted using representative hits (February 2020)^44^. Level 1 gene categories with average frequencies below 0.5 % were omitted. We note that these default settings are permissive and were retained to maintain a common analytical framework across viromes and continuity with prior analyses. Read-level annotation was therefore used here for relative comparisons of broad functional representation among identically processed viromes, rather than to establish the function of any individual read. Because these gene frequencies are proportions of the total annotated genes within each virome, the data are compositional: increases in the relative frequency of one category necessarily entail decreases in others. We therefore interpret these values as relative community composition rather than absolute gene abundance.

### Pathway annotation

Pathway visualizations were guided by KEGG maps (www.genome.jp/), pathway-specific Wikipedia pages (www.wikipedia.org/), and enzyme Wikigenes entries (www.wikigenes.org/). Pathway maps were drawn using Enzyme Commission (EC) Number names of each enzyme (**Table 1**).

**Table 1:**
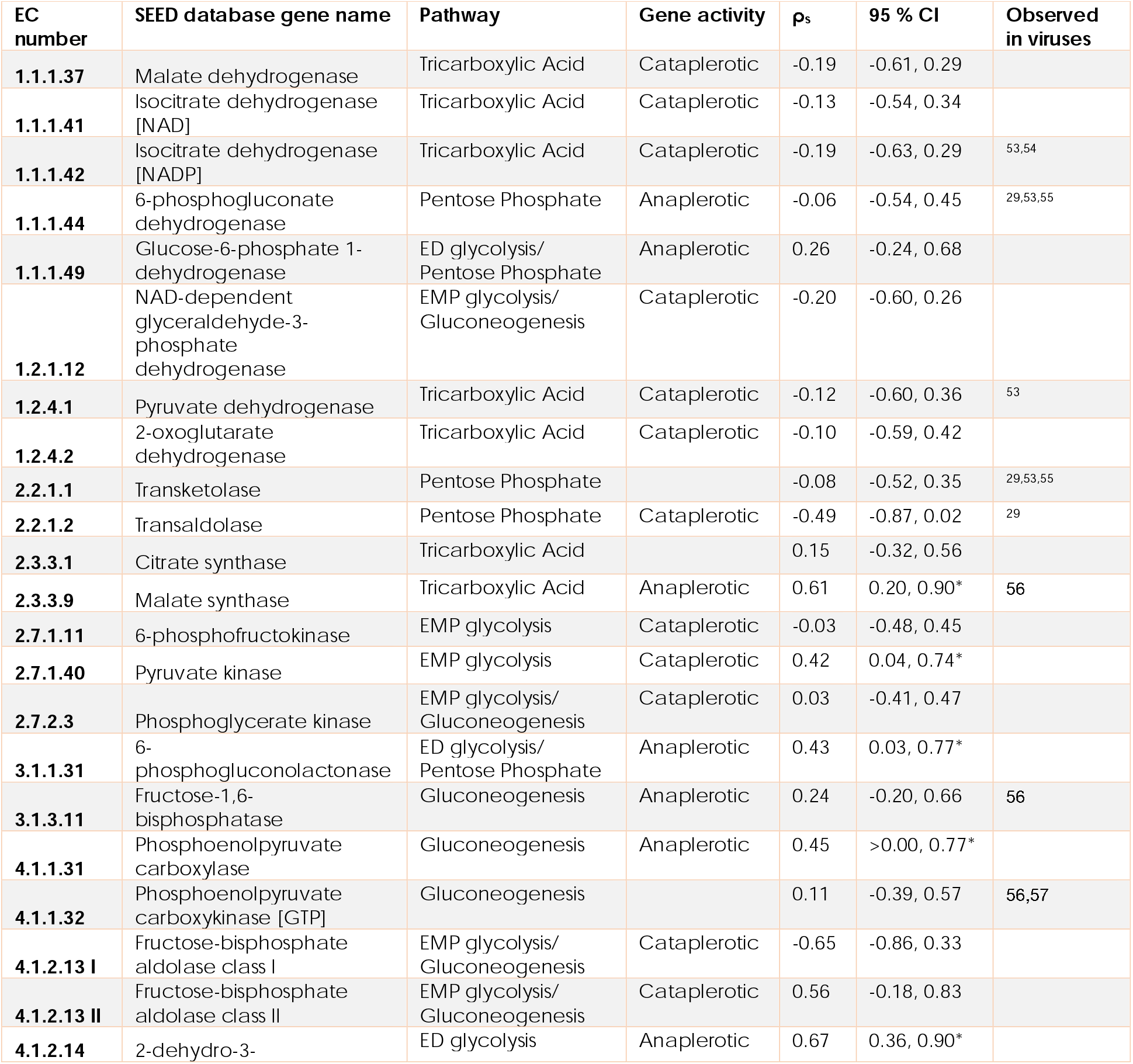
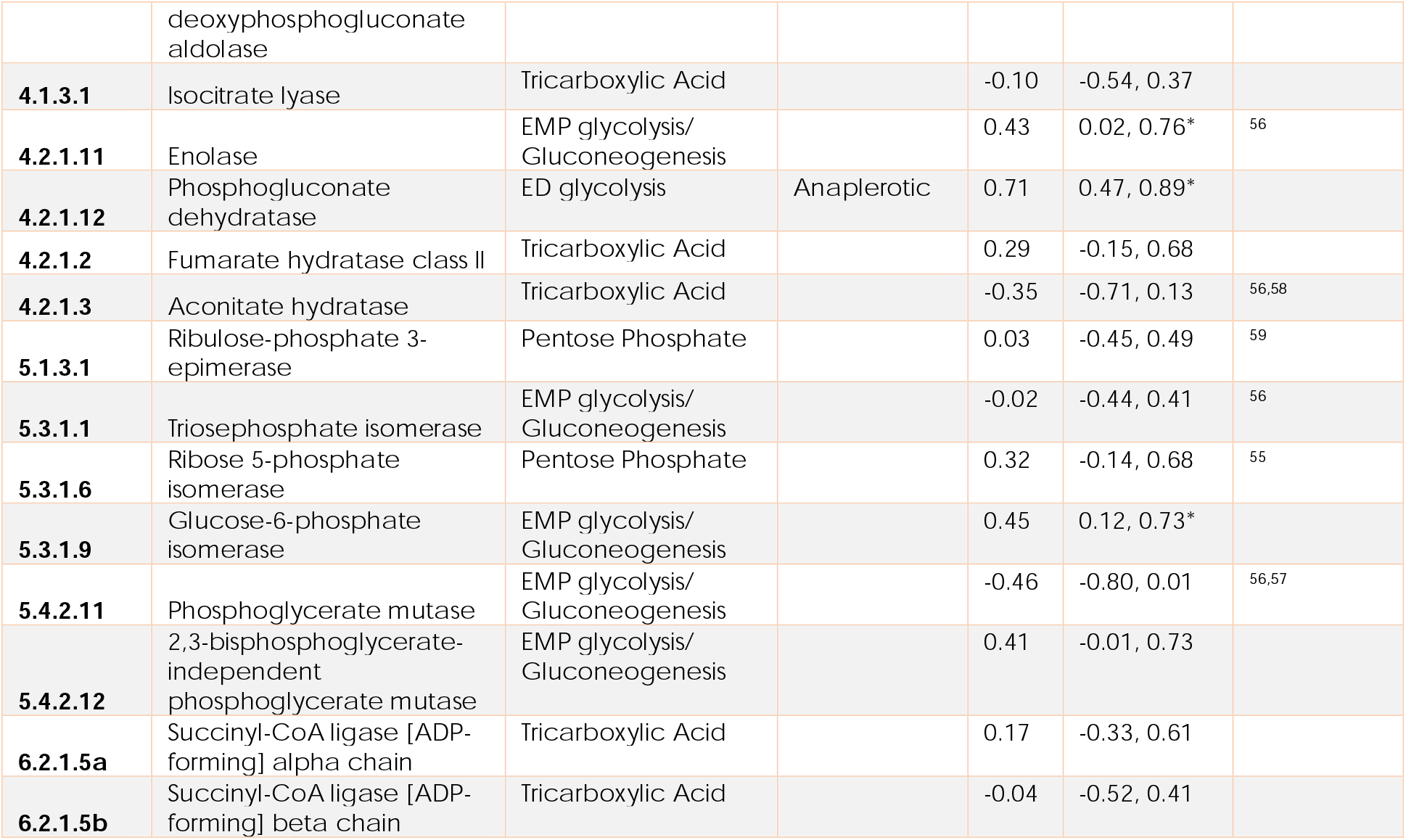
Central Carbon Metabolism genes shown in Figures 2, 3, and 4. Genes are categorized by pathway and gene function, identified by EC number and gene name, and annotated with prior references if they have been observed in viruses before. The correlation between each gene’s relative frequency and viral community temperateness is reported as the bootstrapped median ρ_s_ with its 95% confidence interval from 1,000 iterations. Please refer to **Supplementary Figure 2** for bootstrap ρ_s_ distributions. Genes whose 95 % confidence intervals exclude zero are indicated with asterisks; they are provided as a visual guide and do not represent multiple-testing-adjusted significance. Genes shared between pathways are shown with a slash. Note, although anaplerotic and cataplerotic reactions often refer to steps in the TCA, we are using the term more generally here to refer respectively to the generation of central intermediate metabolites (‘filling up’; anaplerotic) and the consumption of central metabolites to generate specialized precursor or cellular energy pools (‘drawing down’; cataplerotic) more generally^33^. Bootstrapping code can be found at https://github.com/hopefulmonstersucla/viral-metabolism/tree/main/code.

### Anaplerotic and cataplerotic gene categorization

Genes were categorized based on whether they are best known to (i) generate central metabolite pools like glyceraldehyde-3-phosphate (G3P) and pyruvate (‘anaplerotic’ or ‘filling up’ reactions) that are flexibly used in multiple pathways by cells, or (ii) use these central metabolite pools to generate cellular energy or metabolic precursor products that are used for specific purposes (i.e., the products are ‘used’ metabolically; known together as ‘cataplerotic’ or ‘drawing down’ reactions), or (iii) were neither anaplerotic nor cataplerotic (‘neither’). Note that although anaplerotic and cataplerotic reactions often refer to steps in the TCA, we are using the term more generally here (see **Figure 3** for enzyme directionality)^33^.

**Figure 1:**
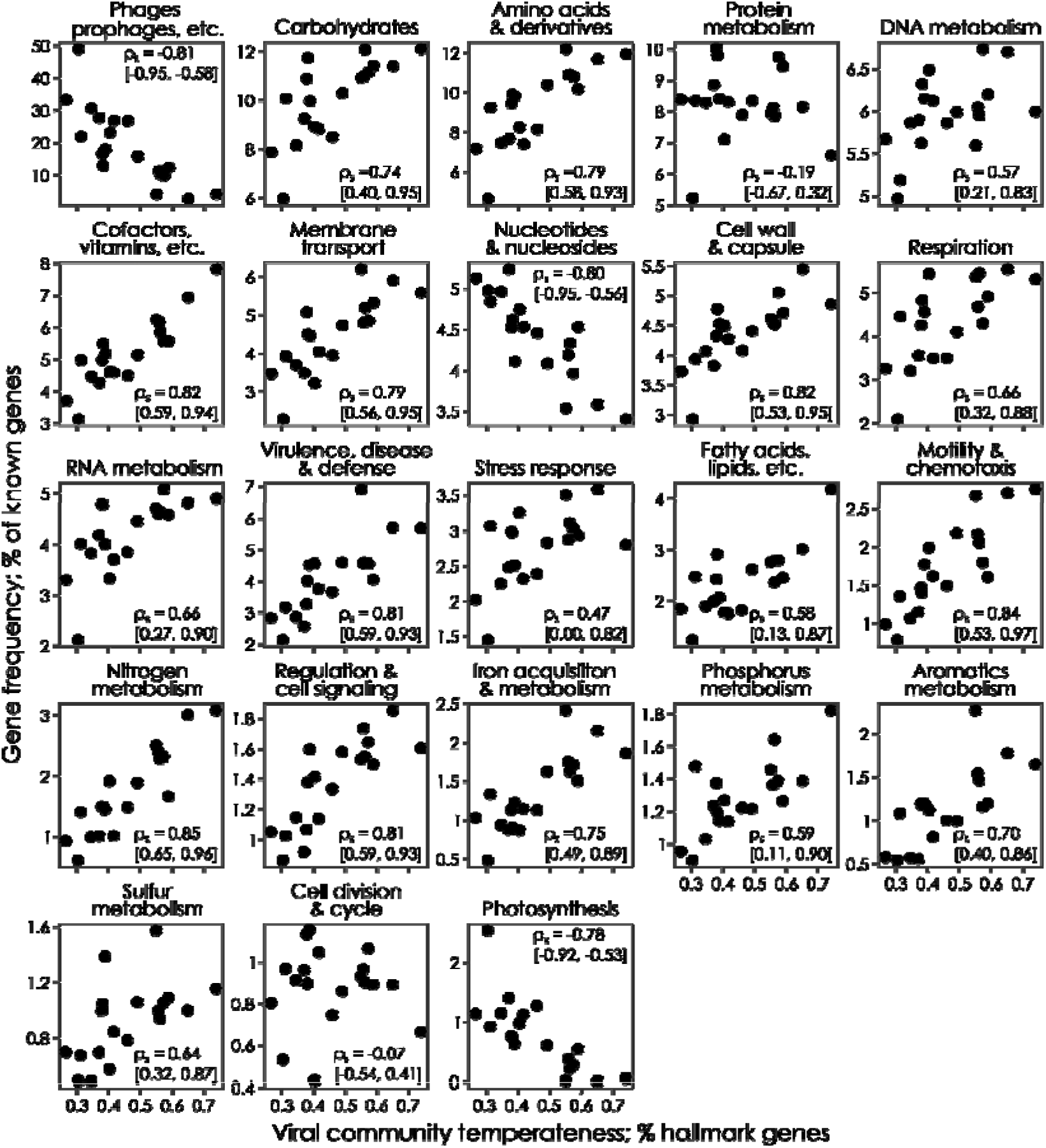
Correlations between metabolic function and community temperateness across the breadth of cellular metabolism. Spearman’s rho estimates (ρ□) are shown with 95% confidence intervals (CIs) in square brackets from 1,000 bootstrapped simulations. Positive Spearman’s ρ values indicate temperate-associated gene categories, whereas negative ρ values indicate lytic-associated gene categories. Panels are ordered by mean gene frequency. The y-axis shows gene frequency as a percentage of known genes, an internally normalized and relative metric for which the frequencies for a given sample across all panels sum to 100%. Gene categories with average frequencies below 0.5% were omitted. Viral community temperateness was calculated as the percentage of integrase and excision genes among the known genes in each virome. The 95% confidence intervals quantify uncertainty and do not represent multiple-testing-adjusted significance. Bootstrapping code can be found at https://github.com/hopefulmonstersucla/viral-metabolism/tree/main/code.

**Figure 2:**
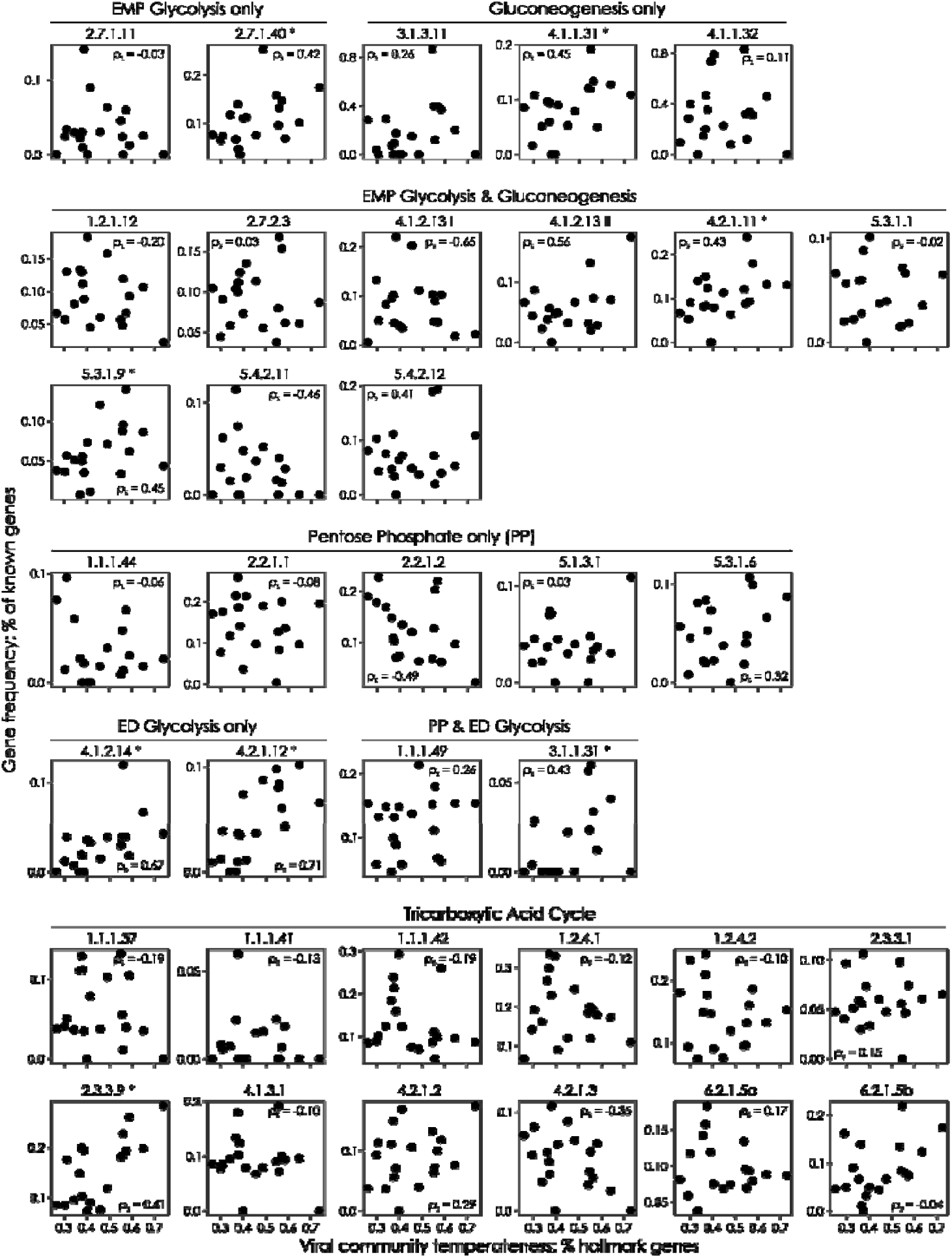
Correlation between central carbon metabolism genes and viral community temperateness. Spearman’s rho estimates (ρ° ) are shown from 1,000 bootstrapped simulations correlating central carbon metabolism gene frequency as a percentage of known genes vs viral community temperateness. Positive Spearman’s ρ values indicate temperate-associated gene categories, whereas negative ρ values indicate lytic-associated gene categories. Panels are ordered by pathway, including Embden-Meyerhof-Parnas glycolysis (EMP), Entner-Doudoroff glycolysis (ED), pentose phosphate (PP), and tricarboxylic acid cycle pathways. Viral community temperateness was calculated as the percentage of integrase and excision genes in the known genes in each virome. Asterisks indicate enzymes whose 95 % confidence intervals exclude zero; they are provided as a visual guide and do not represent multiple-testing-adjusted significance. Genes are labeled by EC numbers; see **Table 1** for gene names and summary statistics.

**Figure 3:**
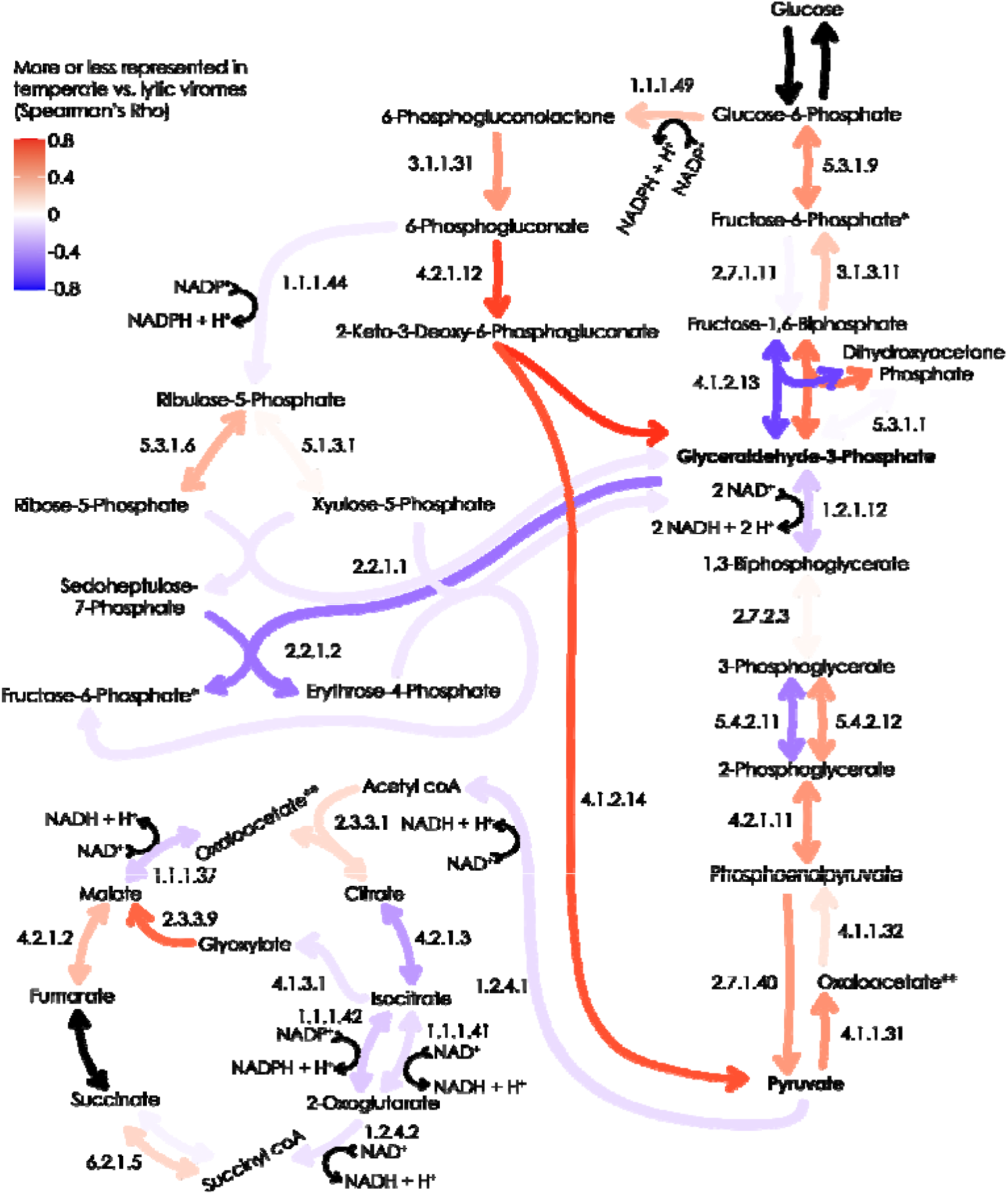
Pathway map of changes in the frequency of central carbon metabolism genes across the lytic to temperate spectrum. Map of central carbon metabolism with each enzymatic step colored by the change in enzymatic metagenomic gene frequency across lytic to temperate viral communities. Enzymes are labeled by EC numbers (see **Table 1** for gene names). Red arrows show higher frequency in temperate communities while blue arrows are lytic associated, reflecting ρ° values for each enzyme as seen in **Table 1** and **Figure 2**. Single-headed and double-headed arrows represent irreversible and reversible reactions, respectively. Black arrows show genes that were absent from all viromes. Central intermediate metabolites glyceraldehyde-3-phosphate (G3P) and pyruvate are shown in bold. *Note that fructose-6-phosphate is found in the EMP/gluconeogenesis pathways as well as the PPP, ** that oxaloacetate is produced in both the TCA and gluconeogenesis, and that EC1.1.1.42 in the TCA has an intermediate oxalosuccinate product/substrate. Code can be found at https://github.com/hopefulmonstersucla/viral-metabolism/tree/main/code.

### Statistical analyses

The correlation between gene frequency, either at a Level 1 SEED category or individual gene level, and viral community temperateness was assessed using Spearman’s Rho (ρ_s_), a non-parametric rank-based measure of how monotonic the relationship between x- and y-values is (i.e., it is a non-parametric correlation coefficient). For supplementary diversity and photosynthesis gene analyses we estimated ρ_s_ using the R cor.test() function with exact = FALSE to avoid ties. For our main analyses shown in **Table 1** and in **Figures 1, 2, 3**, and **4**, we estimated ρ_s_ values using the scipy.stats.spearmanr() function in the Python scipy.stats package with bootstrapping to ensure robust median ρ_s_ assessment over 1000 iterations of bootstrapping with replacement (see **Supplementary Figure 2** for resulting distributions for central carbon metabolism genes). Confidence intervals of the median were assessed as the values corresponding to the 2.5th and 97.5th percentiles of the median values obtained from bootstrapping (95 % CI). For clarity, 95 % confidence intervals are listed in square brackets, as ρ_s_ [95 % CI minimum, 95 % CI maximum]. Confidence intervals are presented to quantify uncertainty around each bootstrapped median estimate. Bootstrapping code can be found at https://github.com/hopefulmonstersucla/viral-metabolism/tree/main/code.

Shannon diversity (H’) was calculated from the relative frequencies of annotated functional genes (“known genes”) within each viral metagenome. Diversity metrics were assessed using the R vegan package, with the diversity() command used for Shannon diversity (natural log-transformed; H’). Median ρ_s_ at the pathway level were calculated as the median value of all of the ρ_s_ values for each enzyme in the pathway (i.e., they are a simple overall median).

### Analytical framework

Viral community temperateness was quantified as the frequency of genes annotated to the SEED Level 3 category Phage_integration_and_excision, following our previous use of integration and excision genes as a hallmark, community-level proxy for temperateness^14^. Gene frequency was calculated as the number of hits to a given gene divided by the total number of annotated (“known”) genes within that metagenome (i.e., *percent known*)^14^. Higher-order functional categories (e.g., the SEED Level 1 categories shown in **Figure 1**) were calculated by summing the frequencies of their constituent genes. This within-sample normalization allowed gene frequencies to be compared across metagenomes with differing sequencing depths. For each metabolic gene or pathway, we calculated Spearman’s rho (ρ_s_) between its frequency and viral community temperateness across the 19 viromes, as shown in **Figures 1-4**. Positive ρ_s_ values indicate genes occurring at higher relative frequencies across the viral community temperateness gradient, whereas negative values indicate enrichment toward the lytic end of the gradient. Gene names, pathways, EC numbers, median ρ_s_ values, 95% confidence intervals, biochemical classifications, and previous reports of these genes in viruses are summarized in **Table 1. Figure 5** then uses these observations, together with established ecological knowledge from prior work, to develop a qualitative conceptual model that explores their possible ecological implications.

### Qualitative conceptual model

We created a qualitative conceptual model to explore how the observed metabolic shifts identified here might interact with the established feedback loops underlying reef microbialization (**Figure 5**). The model is qualitative rather than quantitative and was designed to aid understanding of how viral metabolic capacity may influence ecosystem dynamics when incorporated into existing ecological feedbacks. Parameter values were implemented as relative high and low values to reflect established differences between healthy and degraded reefs (**Supplementary Table 2** for parameter values). For example, high rates of disease^3,45,46^ and bacterial metabolism^1,9^ were used in degraded coral reef simulations, while high lysis rates were used in healthy systems^14,17^.

The model is intended to explore how the observed viral metabolic shifts might alter established ecological feedbacks, rather than to quantitatively predict reef dynamics. The model was initialized with Bacterial, viral, coral, and algae populations at 0.5 arbitrary units. The model was implemented using an iterative loop over six iterations (i.e., time in arbitrary units). This number of iterations was chosen as a balance of having enough iterations to see qualitative population trends, but without the need to make a more complicated model to generate long-term dynamics. At the beginning of each time step, new abundance values for each species (B_new_, C_new_, V_new_) except algae were calculated based on the values from the previous iteration (A_new_). These updated values were applied at the end of each time step. Algae abundance was calculated last, occupying any remaining share of the benthos not occupied by coral. Once a population reached or fell below zero, it was excluded from further dynamics and not allowed to re-enter the ecosystem. Code can be found at https://github.com/hopefulmonstersucla/viral-metabolism/tree/main/code.

For the ‘no viruses’ model shown in **Figures 5a** and **5d**, we used the following equations, where terms with changed parameter values between **Figures 5a** and **5d** are in bold (see **Supplementary Table 2**):

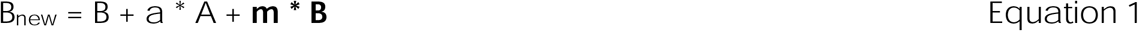

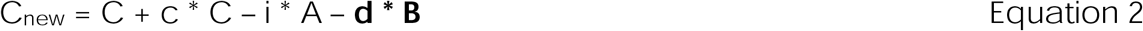

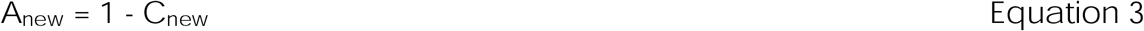

In **Equation 1**, bacteria (B) proliferate due to carbon provided by algae, with carbon supply itself a product of algal population (A) as well as the rate of delivery (a). To make the model as conservative as possible, we did not vary the rate of carbon delivered to bacteria by the algae, although it likely is higher on degraded reefs and enhances bacterial growth more than shown by the model^10^. Bacterial proliferation is also enhanced by metabolic changes (m) reinforced by bacterial population size on degraded reefs^1^. Note that, in the absence of viruses the bacteria have no loss processes, including natural mortality, even though their proliferation may be constrained by a lack of algae to support growth.

In **Equation 2**, coral populations (C) proliferate as a product of coral population size and growth rate (c). Coral growth is limited by algal population size and allelochemical inhibition (i)^8^ as well as by bacterial pathogenicity causing coral disease (d)^3,46^. To make the model as conservative as possible, we did not alter the direct inhibitory effect of algae on coral across healthy and degraded reefs, although it is enhanced on unhealthy reefs^3,7,8,44,46^.

In **Equation 3**, algal population size (A) is solved in the last step of each iteration as the remainder of the benthos not occupied by coral (1 – C) to account for the fact that the benthos is a finite space.

Note that **Equations 2** and **3** are used for all iterations of the model with or without viral lysis and metabolism.

For the ‘with lytic viruses’ model shown in **Figures 5b** and **5e** we used the following equations, with bold terms reflecting varied parameter values and with new terms compared to **Equations 1** and **Equation 2** underlined:

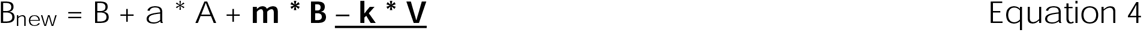

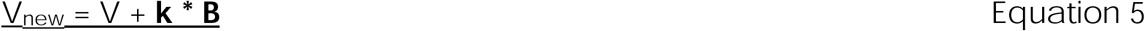

Building on **Equation 1**, lytic viruses are introduced in **Equation 4** as a mortality term for bacteria as an outcome of viral population size (V) and lysis rate (k).

A viral population (V) is added to the model in **Equation 5**, where viruses proliferate at lysis rate (k) from lysis of bacteria. Note that k changes between healthy and degraded reef scenarios, consistent with Piggyback-the-Winner dynamics suppressing lysis on degraded reefs^14^.

For the ‘with viral lysis and metabolism’ model shown in **Figures 5c** and **5f** we used the following equations, with bold terms reflecting varied parameter values and with new terms compared to **Equations 4** and **5** underlined:

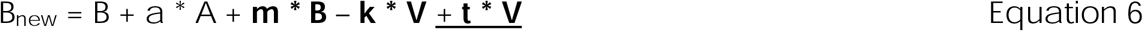

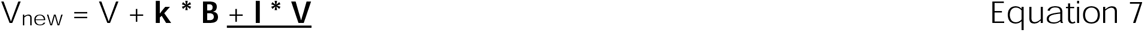

This model builds on **Equation 4** by adding viral metabolic function (t) that enhances bacterial proliferation in degraded reef scenarios associated with temperate Piggyback-the-Winner dynamics in **Equation 6**^14^. **Equation 7** builds on **Equation 5** by adding lytic functions that enhance virion production in healthy reefs (l).

## Results

### Temperate viral communities are enriched in metabolic genes

The frequency of genes in the Phages, Prophages, Transposable Elements, and Plasmids category declined with increasing temperateness of viral communities (Figure 1; bootstrapped median ρ_s_ = −0.81, 95% confidence interval [−0.95, −0.58]; note that medians and confidence intervals are shown as bootstrapped median [95 % confidence interval of the median] throughout the paper). In contrast, the frequency and diversity of ‘bacterial’ metabolic genes that encode known cellular functions increased in temperate viral communities (**Supplementary Figure 1**; ρ_s_ = 0.63 and 0.81, respectively; c.f., genetic diversity declines with temperateness when unknown genes are included^14^). Altogether, this analysis indicates that temperate viral communities are more metabolically similar to bacteria than are lytic communities.

### Photosynthesis genes in lytic communities

The decline in ‘Phages, Prophages, Transposable Elements, and Plasmid’ genes from ∼ 50 % of the known genes in lytic viromes to ∼ 5 % in temperate viromes caused the gene frequency in almost all other categories to increase (as total gene frequencies sum to 100 %; 17 of 22 categories had ρ_s_ > 0.5; **Figure 1**). A notable exception is Photosynthesis genes: these had higher frequencies in lytic communities vs. temperate communities in the Level 1 analysis (ρ_s_ = -0.78 [-0.92, -0.53]; **Figure 1**). Photosynthesis genes, including rhodopsins, were also lytic-associated at the individual-gene level, where 16 out of 20 photosynthesis genes had negative ρ_s_ values (**Supplementary Table 3**). Finally, photosynthesis-related electron transport (Pet) genes that couple Photosystems II and I are absent from the viromes (**Supplementary Table 3**).

### Carbohydrate metabolism genes

Given the high prevalence of carbohydrate genes in the viromes, their elevated frequency in temperate communities (**Figure 1;** ρ_s_ > 0), and their ecological importance, we next focused on genes comprising central carbon metabolism^1,24,25^: Embden-Meyerhof-Parnas (EMP) and Entner-Doudoroff (ED) glycolysis pathways, the pentose phosphate pathway (PPP) and the Tricarboxylic Acid Cycle (TCA; **Figure 2**). Note that for this and subsequent analyses we retained only sequences with hits to SEED database entries with high-confidence Enzyme Commission (EC) annotations^47^.

### Entner-Doudoroff genes are enriched in temperate viromes

Few patterns were readily apparent from the ρ_s_ values for genes in each pathway (**Figure 2; Supplementary Figure 2**). When viewed as a pathway map, though, the ED pathway was more clearly enriched in temperate viral communities (i.e., the red arrows running through the center of **Figure 3**). This map identified the ED pathway as the only pathway that changed consistently across the lytic-temperate spectrum (**Figure 4a**), and was exclusively over-represented in temperate viral communities (i.e., ρ_s_ > 0; EC 1.1.1.49 ρ_s_ = 0.26 [-0.24, 0.68], EC3.1.1.31 ρ_s_ = 0.43 [0.03, 0.77], EC4.1.2.14 ρ_s_ = 0.67 [0.36, 0.90], and EC4.2.1.12 ρ_s_ = 0.71 [0.47, 0.89]). Altogether, this gave the ED pathway an overall median ρ_s_ = 0.55 in contrast to overall median ρ_s_ = 0.03 for EMP glycolysis, ρ_s_ = 0.175 for gluconeogenesis (GNG), ρ_s_ = 0.03 for the PPP, and ρ_s_ = -0.10 for the TCA (see **Figure 4a**).

**Figure 4:**
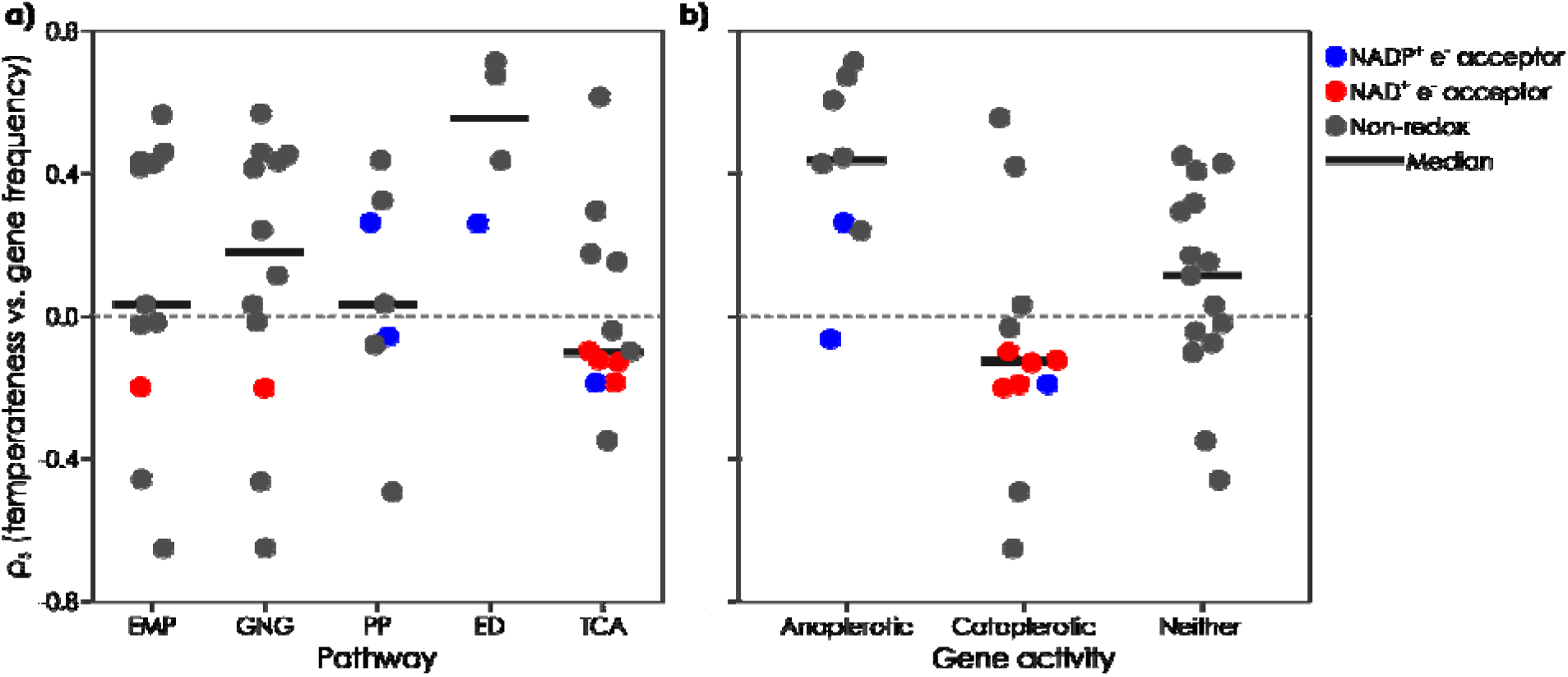
Correlations between viral community temperateness and central carbon gene frequency across pathways and gene function. **(a)** Spearman’s rho correlation coefficients (ρ° ) for genes in the Embden-Meyerhof-Parnas glycolysis (EMP), gluconeogenesis (GNG), pentose phosphate (PP), Entner-Doudoroff glycolysis (ED), and tricarboxylic acid cycle (TCA) pathways are shown with median values marked by horizontal bars in each pathway. (b) Coefficients are also shown for genes involved in building up central intermediate metabolite pools like pyruvate and glyceraldehyde-3-phosphate (anaplerotic reactions) or consuming general metabolites to make specialized pools of precursors and cellular energy (cataplerotic reactions) in central carbon metabolism, compared with genes involved in neither anaplerosis nor cataplerosis, also with median values shown. Oxidoreductase genes are colored by their electron acceptor: blue for NADP^+^ electron acceptors involved in biomass generation, and red for NAD^+^ electron acceptors involved in cellular energy generation. Dashed horizontal lines indicate ρ° = 0. Note, although anaplerotic and cataplerotic reactions often refer to steps in the TCA, we are using the term more generally here to refer respectively to the generation (‘filling up’; anaplerotic) of flexible general intermediate metabolites and the consumption of general metabolites towards precursor and cellular energy production (‘drawing down’; cataplerotic) more generally^33^.

**Figure 5:**
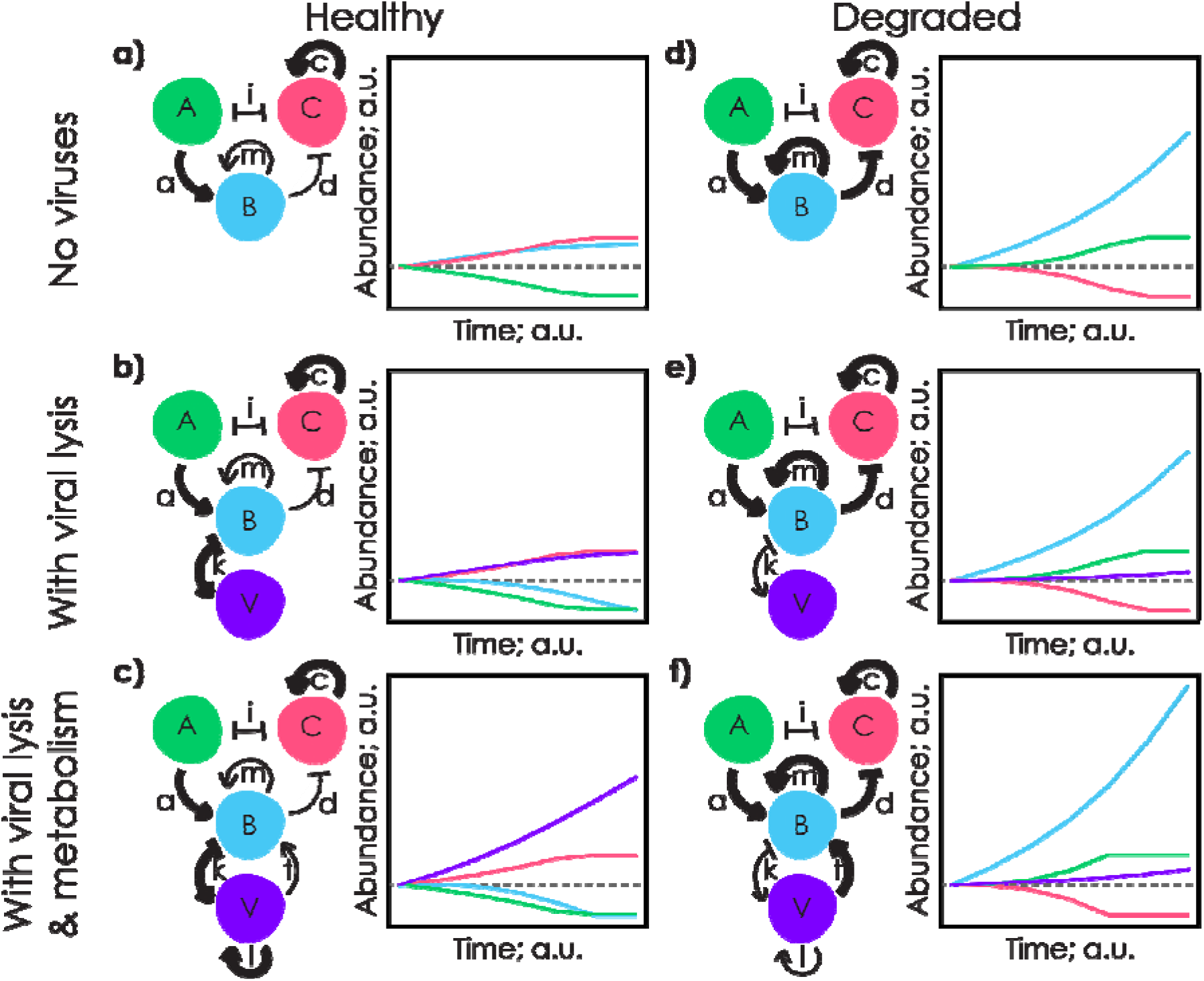
Conceptual model illustrating predicted ecosystem consequences of lytic- and temperate-associated viral metabolism. The predicted effects of viral lysis and metabolic capability on algal (A; green), coral (C; coral), bacteria (B; blue) and viral (V; violet) population sizes in healthy and degraded reef scenarios with no viruses (a and d, respectively), with viral lysis (b and e, respectively), and with viral lysis and metabolism (c and f, respectively). Compartment models showing how the model was implemented for each panel are shown (left) as well as resulting dynamics (right). Dynamics show changes in populations over time, in arbitrary units (a.u.), reflecting the qualitative nature of the model. In each iteration, algal population size A was solved as the 1 – C to reflect the finite space available on the benthos. Parameter values, shown as line weights, qualitatively reflect known processes on healthy vs degraded reefs, and can be found in **Supplementary Table 2**. Rates include: (i) coral-algal inhibition, (c) coral proliferation, (a) algal-mediated DOC supply to bacteria, (m) bacterial anabolic metabolism, (d) bacterial-mediated coral disease, (k) viral lysis of bacteria, (l) lytic viral metabolism, and (t) temperate viral metabolism. Note that panels **(b)** and **(e)** depict the Kill-the-Winner and Piggyback-the-Winner models, respectively. Algal, coral, bacteria, and viral pools were initialized at 0.5 for all simulations where present. To make the model as conservative as possible, rates of algal carbon delivery (a), and direct coral-algal interference (i) were not changed across scenarios; doing so would lead to enhanced loss of coral. Note that flat lines at minimal values reflect functional population extinction. Code can be found at https://github.com/hopefulmonstersucla/viral-metabolism/tree/main/code.

### Anaplerotic temperate metabolism

Given the role of ED glycolysis in generating central metabolite pools for use in diverse pathways (i.e., ED is an anaplerotic pathway), we investigated whether temperate viral communities were enriched in anaplerotic genes more generally (see **Table 1** for a list of anaplerotic and cataplerotic genes)^33^. Temperate viral communities were, indeed, enriched in anaplerotic genes, with an overall median ρ_s_ = 0.44 for anaplerotic reactions and only one gene even weakly lytic associated (6-phosphogluconate dehydrogenase EC 1.1.1.44 from the PPP; ρ_s_ = -0.06 [-0.54, 0.45]; **Figure 4b**).

### Amino acid precursor genes are enriched in lytic viromes

In contrast to the ED-enrichment of temperate communities, the pathway map does not show many areas of contiguous lytic-associated genes (**Figure 3**). However, several non-oxidative PPP genes were enriched in lytic communities, especially transaldolase EC 2.2.1.2 (ρ_s_ = -0.49 [-0.87, 0.02]). This enzyme consumes the central metabolite pool of glyceraldehyde-3-phosphate (G3P) to produce fructose 6-phosphate and erythrose 4-phosphate. Given that other fructose-6-phosphate producing genes are not similarly enriched in lytic communities (e.g., EC 2.2.1.1 in the PPP ρ_s_ = -0.08 [-0.52, 0.35], EC5.3.1.9 in EMP ρ_s_ = 0.45 [0.12, 0.73], and EC3.1.3.11 in gluconeogenesis ρ_s_ = 0.24 [-0.20, 0.66]), this suggests that the primary outcome of the non-oxidative PPP in lytic viral communities is the cataplerotic consumption of G3P for the generation of erythrose-4-phosphate, a precursor for amino acid biosynthesis^48^.

### Redox genes are over-represented in lytic communities

Given this cataplerotic focus in lytic communities, we investigated whether lytic communities were more strongly associated with individual cataplerotic reactions regardless of pathway rather than with whole pathways. Focusing on the distribution of anabolic redox steps across all the central carbon metabolism pathways, we found that NADPH-generating reactions were not consistently associated with either lytic or temperate communities (EC1.1.1.44 ρ_s_ = -0.06, EC1.1.1.42 ρ_s_ = -0.19, and EC1.1.1.49 ρ_s_ = 0.26, giving a median ρ_s_ = -0.06). In contrast, NADH-producing oxidoreductase genes were universally over-represented in lytic communities (i.e., all ρ_s_ < 0; median ρ_s_ = -0.13 across all NADH-producing reactions from EC1.1.1.37 ρ_s_ = -0.19 [-0.61, 0.29], EC1.1.1.41 ρ_s_ = -0.13 [-0.54, 0.34], EC1.2.1.12 ρ_s_ = -0.20 [-0.60, 0.26], EC1.2.4.1 ρ_s_ = -0.12 [-0.60, 0.36], EC1.2.4.2 ρ_s_ = - 0.10 [-0.59, 0.42]; **Table 1**; **Figures 2, 3**, and **4**). This included all redox genes in the TCA being lytic associated (i.e., ρ_s_ < 0, with an overall median ρ_s_ = -0.13) in contrast to the other, non-oxidoreductase genes which had a median ρ_s_ = 0.15, highlighting the specific overrepresentation of redox genes in lytic communities (**Figure 4a**).

### Cataplerotic lytic metabolism

Given the association between lytic communities and cataplerotic reactions observed in the PPP and the TCA, we then investigated whether lytic communities are broadly enriched in cataplerotic genes. We observed that the overall median ρ_s_ of cataplerotic reactions was - 0.125 (i.e., these genes were enriched in lytic communities) compared to ρ_s_ = 0.44 and ρ_s_ = 0.11 for anaplerotic reactions and reactions that are neither cataplerotic nor anaplerotic, respectively. This result highlights the association between lytic viral communities and cataplerotic metabolism.

### Conceptual Model Predictions for Healthy Coral Reefs

To explore the potential ecological implications of the observed metabolic shifts, we developed a qualitative conceptual model of algal-bacterial-viral-coral interactions, with all population sizes measured in illustrative arbitrary units (**Figure 5**). Parameterizing the model to healthy reef values (**Figure 5a**; note that parameterization of the healthy reef scenario largely insulates coral from the effects of bacterial overgrowth due to the small d value used; see **Supplementary Table 2** for parameters and values)^1,3,9,14,45^, the system became coral-dominated by t = 5, with coral-bacterial coexistence (coral and bacteria had increased from 0.5 a.u. initial values to 1.0 a.u. and 0.89 a.u., respectively, by t = 6). While the addition of viral lysis (**Figure 5b**) did not markedly change coral or algal dynamics, it led to the extinction of bacteria by t = 6 and to an ecosystem dominated by coral and viruses (coral and viruses had increased from 0.5 a.u. to 1.0 a.u. and 0.97 a.u., respectively, by the end of the simulation). Finally, adding in viral metabolism led to an earlier extinction of bacteria (by t = 5) and enhanced viral population size (viruses were 2.29 a.u. by t = 6), although coral and algal population dynamics were unaffected (**Figure 5c**). Altogether, the conceptual model predicts that incorporating lytic-associated viral metabolism could strengthen bacterial suppression and increase viral populations while preserving the coral-dominated state characteristic of healthy reefs.

### Conceptual Model Predictions for Degraded Coral Reefs

In contrast to healthy systems, our model suggests that degraded coral reefs would be overgrown with algae rather than coral (**Figure 5**). Note that the parameterization of the degraded reef scenarios leads to enhanced deleterious bacterial effects on coral health. Coral went extinct by t = 5 in the degraded scenarios without viruses and with viral lysis only (**Figure 5d and e**), as algae overgrew the benthos and bacteria proliferated to levels much higher than in any of the healthy scenarios (B = 2.75 a.u. and 2.64 a.u. in the no virus and lysis-only models, respectively, by t = 6). In contrast to bacteria, viral populations did not proliferate markedly in the viral-lysis model, increasing from 0.5 a.u. to only 0.64 a.u. by t = 6 (**Figure 5e**). Adding in viral metabolism that enhanced bacterial proliferation (t) allowed bacteria to proliferate even further (reaching 3.80 a.u.), supporting a modest rise in viruses (to 0.75 a.u.), and driving the coral to extinction sooner (by t = 4; **Figure 5f**). Altogether, the conceptual model predicts that viral metabolism supporting bacterial proliferation may enhance microbial overgrowth and promote shifts toward algal dominance of the benthos.

## Discussion

The role of microbial overgrowth in coral reef decline is well established^1–3^. It is further intensified by shifts in bacterial metabolism and viral lifestyles that promote bacterial proliferation^14,17^. Here, we found that lytic- and temperate-associated viral communities also differ systematically in their central carbon metabolic gene repertoires. When interpreted within the framework and assumptions of our conceptual model, these contrasting metabolic profiles suggest that viral metabolism may reinforce either reef resilience or microbialization. Because switching between lytic and temperate infection is closely linked to bacterial physiology and environmental conditions such as nutrient availability^14,35,49,50^, these findings further suggest the hypothesis that manipulating bacterial metabolic state may influence viral infection strategy and metabolism, presenting a potential new avenue for reef conservation.

Viruses become increasingly temperate on degraded reefs, following Piggyback-the-Winner (PtW) dynamics^14^, in which abundant, rapidly proliferating bacterial hosts select for non-lethal, lysogenic infection. This switch in viral lifestyle suspends the top-down control exerted by lytic viruses, allowing bacterial populations to expand unchecked by lethal infection^14,35^. Temperate infection not only reduces lysis, but also promotes horizontal gene transfer of pathogenicity genes^51^, which can help bacteria evade predation by eukaryotic grazers and potentially compromise coral health^14,20–22^. This forms a positive feedback loop wherein bacterial growth triggers viral lifestyle switching, which in turn facilitates further bacterial growth, establishing a ‘Temperate Ratchet’^23^ that locks ecosystems into ever-exacerbating bacterial overgrowth and coral disease. Worse, we found that temperate-associated viral communities are enriched in metabolic genes, including those involved in Entner-Doudoroff glycolysis, that are known to support the generation of central metabolite pools for bacterial growth^1^. When interpreted within our conceptual model, these metabolic shifts may further reinforce the Temperate Ratchet by promoting bacterial proliferation alongside reduced viral lysis^23^.

Healthy reefs, in contrast, appear to be guarded by lytic viruses. Lytic infection rates are driven by virus-host predator-prey encounter frequencies that are the outcome of host and viral abundances^13–16,52^. As a result, as bacterial populations grow, so should lytic infections and lysis, precluding bacterial overgrowth^13,52^. Further, the viral communities on healthy reefs do not harbor the high frequency of pathogenicity genes seen on degraded reefs, meaning that grazing and coral health are not being hindered by viral-encoded genes, which also helps to keep bacterial densities low^3,14,44^. Here, lytic-associated viral communities were enriched in cataplerotic genes that draw down central metabolite pools toward biosynthetic precursors and reducing power. This signature did not arise from the EMP pathway, which characterizes bacterial metabolism on healthy reefs^1^ but showed little association with viral community temperateness here (pathway median ^ρ_s_^ = 0.03). Instead, it was driven by non-oxidative PPP genes such as transaldolase and by NADH-producing oxidoreductases^29,53^. Within our conceptual model, these metabolic features are predicted to support greater virion production, increasing virus-host encounters, infection, and lysis, a process consistent with recent descriptions of reef ‘viralization.’ Within this framework, lytic-associated viral metabolism and reef viralization are predicted to counterbalance the Temperate Ratchet, helping to stabilize healthy reef ecosystems^23,34^.

Our observations and conceptual model together suggest that viral metabolism associated with the lytic-to-temperate shift may reinforce contrasting ecological trajectories on coral reefs. Building on established ecosystem-scale processes such as viral lifestyle switching^14^, the contrasting lytic- and temperate-associated metabolic profiles identified here are predicted to reinforce contrasting ecological feedbacks, contributing to either resilience maintained through metabolically enhanced lytic control of bacteria or microbialization reinforced by temperate infection and metabolism. Because viral infection strategy and viral metabolism are both linked to bacterial physiology and nutrient availability, interventions that limit nutrient inputs could shift viral communities toward lytic infection and its associated metabolism^4,10,12,14,35,52^. Within this conceptual framework, such interventions may promote a ‘re-viralization’ of reef ecosystems, harnessing viral lifestyle and metabolic switching as a potential mechanism to enhance reef restoration and resilience.

## Data Availability

Viral metagenomes can be found on MG-RAST under the accession numbers 4683670.3, 4683674.3, 4683677.3, 4683680.3, 4683683.3, 4683684.3, 4683686.3, 4683703.3, 4683704.3, 4683712.3, 4683718.3, 4683720.3, 4683731.3, 4683733.3, 4683745.3, 4683746.3, 4683747.3, 4574481.3, and 4574475.3 (see **Supplementary Table 1** for library data summaries; www.mg-rast.org). Fasta files for each sample, statistical and modeling code, and annotations and simulations data can be found at https://github.com/hopefulmonstersucla/viral-metabolism.

## Acknowledgments

This work was supported by a BIG Summer stipend from Simons Foundation (awarded to Alexander Hoffmann for CU and NY) as well as by NSF Award OCE-2201645 to BK that provided Hopeful Monsters Summer stipends to JK, MK, GD, NF, EG, AK, and IT. LWK was supported by NSF award OCE-2118617. RAE was supported by an award from the NIH NIDDK RC2DK116713, an award from the Australian Research Council DP220102915, and an award from the Gordon and Betty Moore Foundation number 9871 (Perpetual Viral Origins). JG was supported by NSF Biology Integration Institute award 2119968. ML was supported by an NSF Postdoctoral Research Fellowship in Biology award 2209377. The authors would like to thank the captain, crew, and staff of the *NOAAS Hi’ialakai* (R 334) and National Oceanic and Atmospheric Administration (NOAA) shore support, as well as the members of the Pacific Reef Assessment and Monitoring Program of NOAA’s National Coral Reef Monitoring Program.

## Supplementary Tables and Figures

**Supplementary Table 1:** Summary of the 19 samples analyzed, including sample accession numbers, the total number of base pairs, reads sequenced, and locations of viral metagenomes. Note that all metagenomes have been previously published except Pearl and Hermes Reef Wreck and Caroline Wreck Site samples indicated by an asterisk. Caroline Island is now named Millennium Atoll. Fasta files for all metagenomes can be found at https://github.com/hopefulmonstersucla/viral-metabolism/tree/main/fastas.

| Sample Location | MG-RAST Accession Number | bp count | Read count | Latitude | Longitude |
| --- | --- | --- | --- | --- | --- |
| Millennium Atoll | 4683670.3 | 47624434 | 173813 | -9.9138 | -150.2038 |
| Flint Atoll | 4683674.3 | 7294624 | 26673 | -11.43 | -151.8192 |
| Starbuck | 4683677.3 | 28130343 | 102666 | -5.6384 | -155.8811 |
| Maui | 4683680.3 | 48918703 | 178536 | 20.86447 | -156.13911 |
| Oahu | 4683683.3 | 28708747 | 104972 | 21.4389 | -158 |
| Holoikauaua (Pearl & Hermes Reef) | 4683684.3 | 211441921 | 771690 | 27.8557 | -175.8318 |
| Kauai | 4683686.3 | 14722075 | 53831 | 22.0964 | -159.5261 |
| Starbuck | 4683703.3 | 51673568 | 188592 | -5.6384 | -155.8811 |
| Niihau | 4683704.3 | 17098607 | 62505 | 21.85115 | -160.18327 |
| Molokai | 4683712.3 | 31626104 | 115424 | 21.17481 | -156.8214 |
| Papaopoho (Lisianski) | 4683718.3 | 51289619 | 187189 | 25.98702 | -173.99438 |
| Malden Atoll | 4683720.3 | 68368950 | 249522 | -4.0123 | -154.9178 |
| Lalo (French Frigate Shoals) | 4683731.3 | 12353528 | 45131 | 23.87748 | -166.29115 |
| Hawaii | 4683733.3 | 30728163 | 112147 | 20.00324 | -155.83318 |
| Lalo (French Frigate Shoals) | 4683745.3 | 64763334 | 236363 | 23.6417 | -166.12928 |
| Lana'i | 4683746.3 | 38160746 | 139273 | 20.76167 | -156.8334 |
| Millennium Atoll | 4683747.3 | 71166319 | 259736 | -9.9138 | -150.2038 |
| Holoikauaua (Pearl & Hermes Reef Wreck*) | 4574481.3 | 24339533 | 88980 | 27.8666 | -175.7468 |
| Caroline Wreck Site* | 4574475.3 | 11555298 | 42262 | -9.9957 | -150.2161 |

**Supplementary Table 2:** Parameters and values used in the qualitative ecosystem model. To isolate the effect of bacterial and viral proliferative genes on algal, coral, bacteria, and viral populations, as many parameters as possible were held constant, and only those relating directly to bacterial and viral proliferation and pathogenicity changed (listed in **bold**). All populations, shown with arbitrary units to show relative change in Figure 5, were initialized at 0.5. Note that, to account for finite space on the benthos, algal population size A is solved from coral population size C as A = 1 – C each iteration. Code can be found at https://github.com/hopefulmonstersucla/viral-metabolism/tree/main/code.

| Parameter | Symbol | Healthy reef without viruses | Degraded reef without viruses | Healthy reef with viruses | Degraded reef with viruses | Healthy reef with viruses and viral metabolism | Degraded reef with viruses and viral metabolism |
| --- | --- | --- | --- | --- | --- | --- | --- |
| **Bacterial replication** | m | 0.02 | 0.2 | 0.02 | 0.2 | 0.02 | 0.2 |
| **Bacterial inhibition of coral** | d | 0.02 | 0.2 | 0.02 | 0.2 | 0.02 | 0.2 |
| Coral replication | c | 0.2 | 0.2 | 0.2 | 0.2 | 0.2 | 0.2 |
| Algal inhibition of coral | i | 0.02 | 0.02 | 0.02 | 0.02 | 0.02 | 0.02 |
| Algal support of bacteria | a | 0.2 | 0.2 | 0.2 | 0.2 | 0.2 | 0.2 |
| **Viral replication** | l | - | - | - | - | 0.2 | 0.02 |
| **Viral lysis of bacteria** | k | - | - | 0.2 | 0.02 | 0.2 | 0.02 |
| **Bacterial support of viral replication** | k | - | - | 0.2 | 0.02 | 0.2 | 0.02 |
| **Viral support of bacterial replication** | t | - | - | - | - | 0.02 | 0.2 |
| Bacterial initial population | B | 0.5 | 0.5 | 0.5 | 0.5 | 0.5 | 0.5 |
| Coral initial population | C | 0.5 | 0.5 | 0.5 | 0.5 | 0.5 | 0.5 |
| Algal initial population | A | 0.5 | 0.5 | 0.5 | 0.5 | 0.5 | 0.5 |
| Viral initial population | V | - | - | 0.5 | 0.5 | 0.5 | 0.5 |

**Supplementary Table 3:**
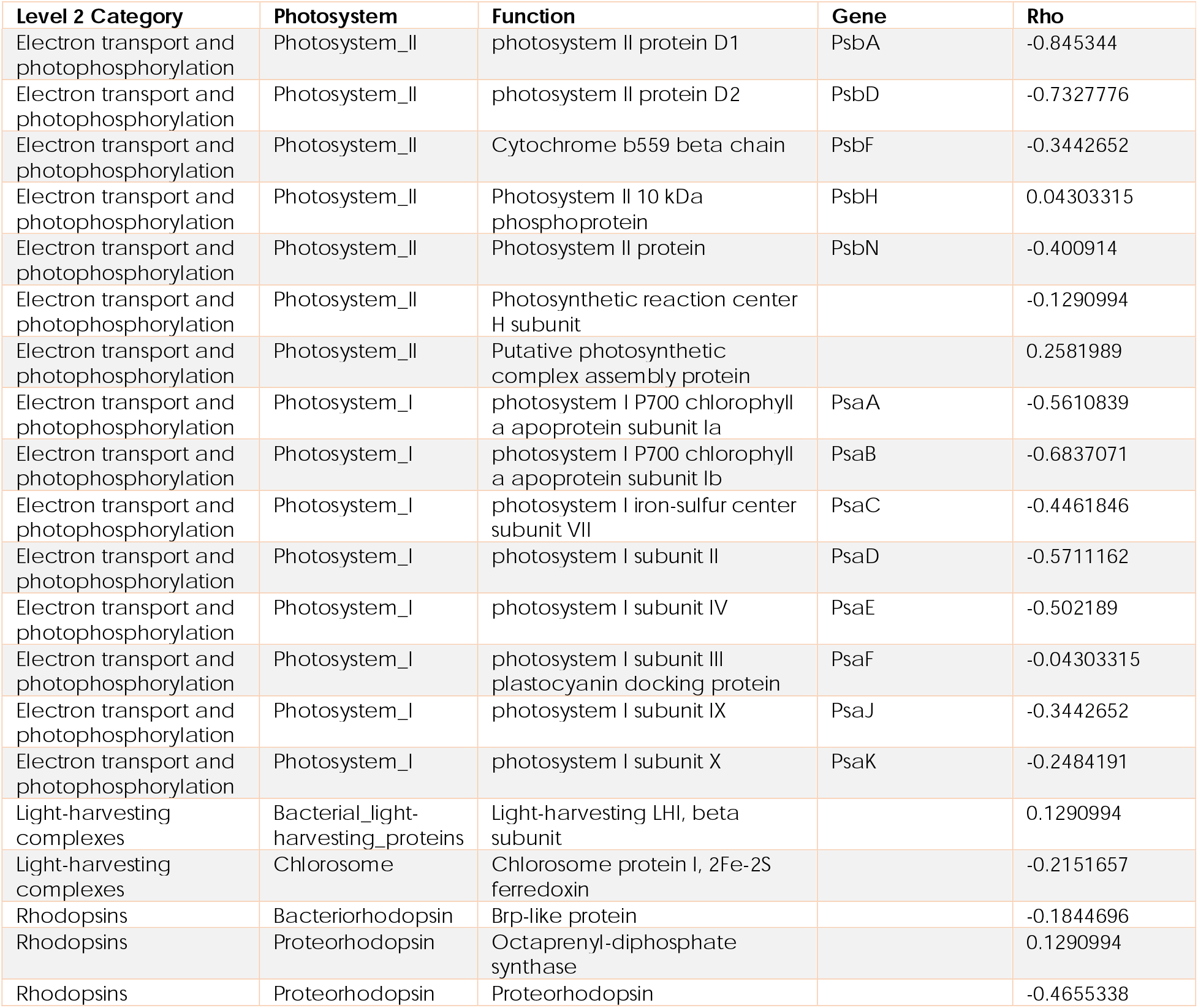
The frequency of photosynthesis genes across the lytic-temperate spectrum. Genes were identified using SEED Level 2 categorization and are organized according to the KEGG photosynthesis map. Negative ρ_s_ values indicate lytic-associated genes and positive ρ_s_ values indicate temperate-associated genes.

**Supplementary Figure 1:**
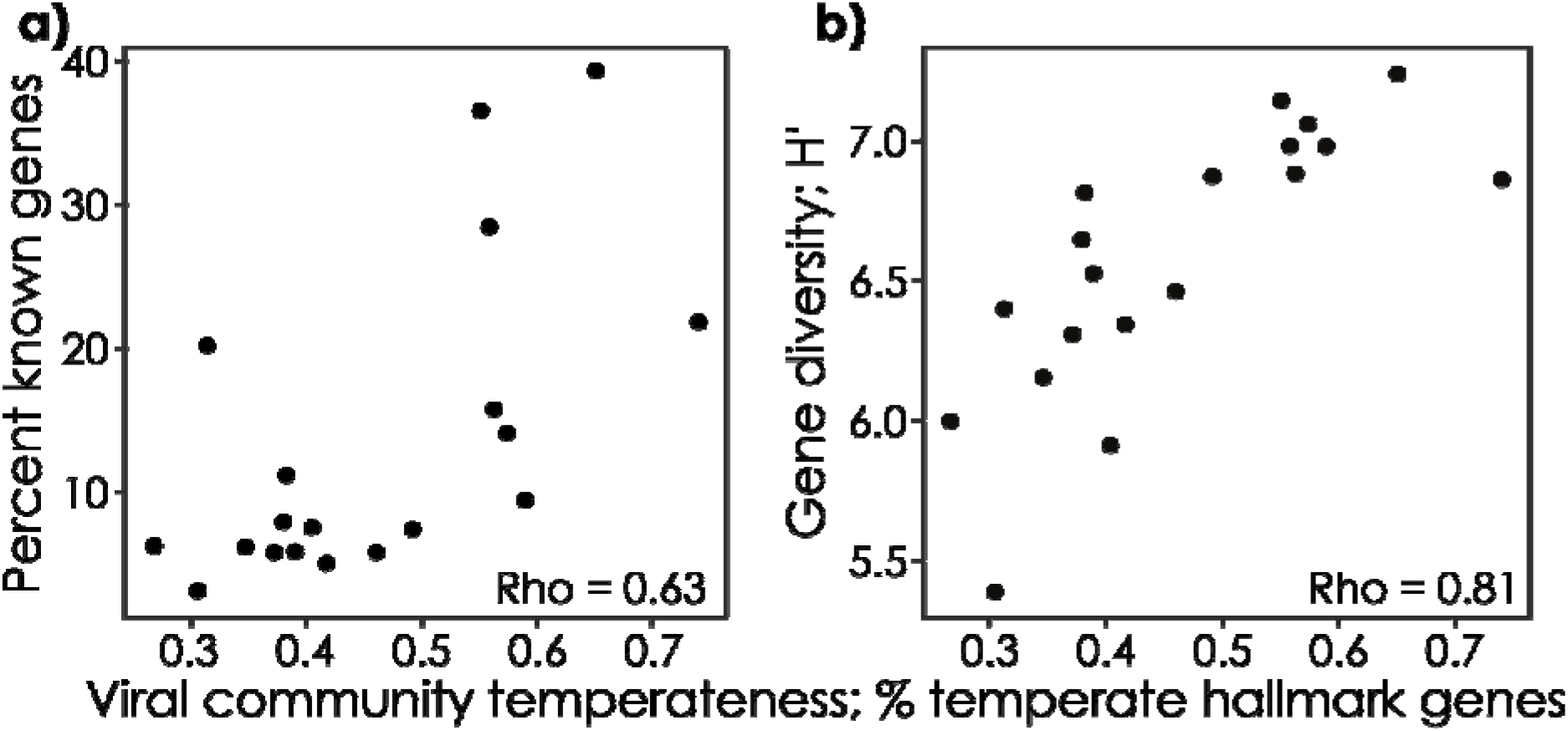
Temperate viral communities are more cell-like and are more functionally diverse in known genes. (a) The percent of genes with hits against the SEED database (i.e., percent known genes), and (b) the diversity of those known genes in the viral communities across the lytic to temperate spectrum. Note that the percent known genes is indicative of how metabolically similar the viruses are to cells given that known genes are commonly identified in the study of cells. Viral community temperateness was calculated as the percentage of integrase and excision genes in the known genes in each virome.

**Supplementary Figure 2:**
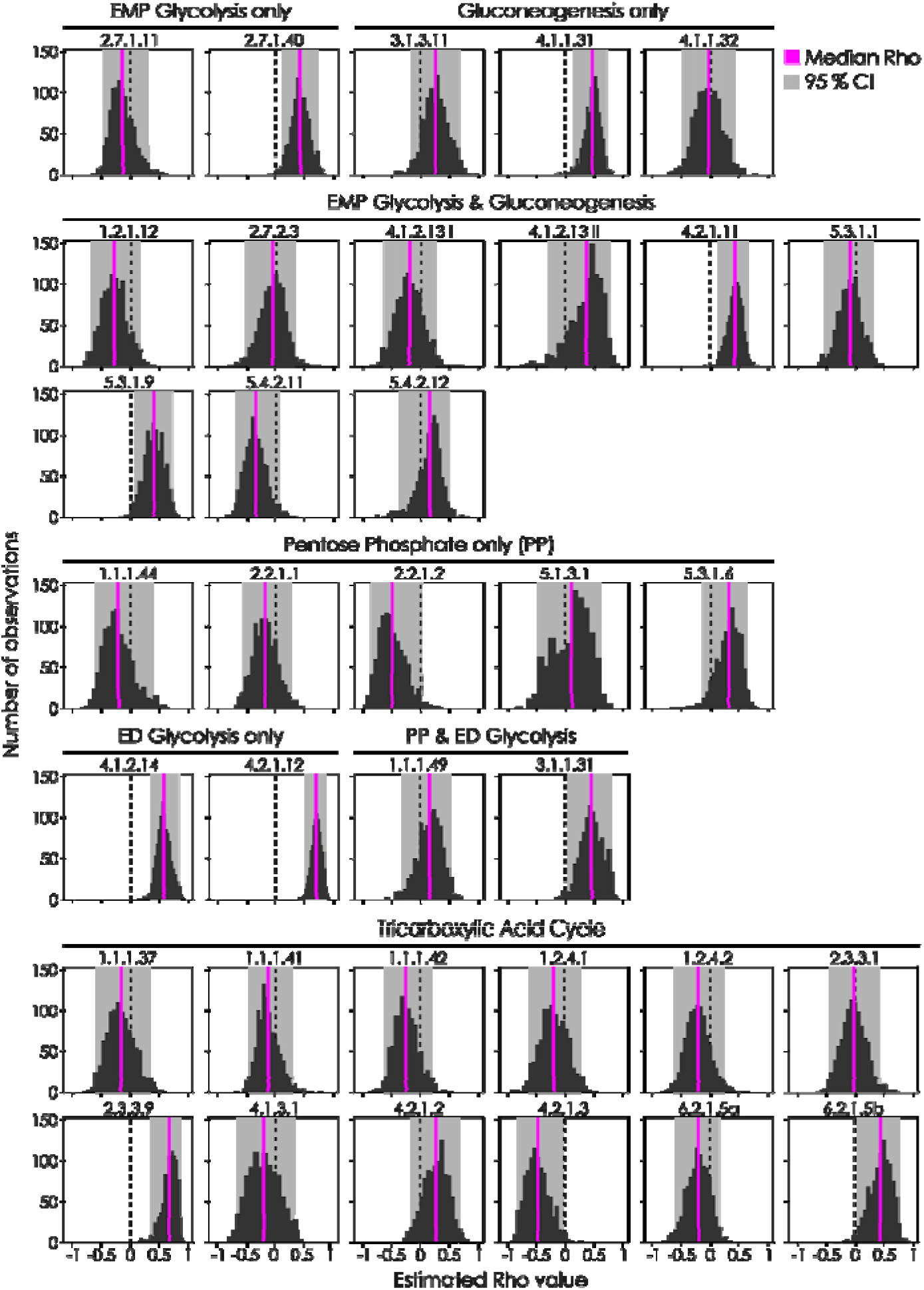
Bootstrapped distribution of ρ_s_ from 1,000 iterations correlating the frequency of central carbon metabolism genes with viral community temperateness (number of observations vs ρ_s_ value). Genes are grouped by pathway. Histogram shows the distribution of bootstrapped ρ_s_ values, with pink lines showing the median bootstrapped value and shaded grey areas showing the 95 % confidence interval of the median ρ_s_. Dashed vertical lines show ρ_s_ = 0. Bootstrapping code can be found at https://github.com/hopefulmonstersucla/viral-metabolism/tree/main/code.

